# Yorkie is required to restrict the injury responses in planarians

**DOI:** 10.1101/092353

**Authors:** Alexander Y.T. Lin, Bret J. Pearson

## Abstract

Regeneration requires the precise integration of cues that initiate proliferation, direct differentiation, and ultimately re-pattern tissues to the proper size and scale. Yet how these processes are integrated with wounding cues remains relatively unknown. The freshwater planarian, *Schmidtea mediterranea*, is an ideal model to study the stereotyped proliferative and transcriptional responses to injury due to its high capacity for regeneration. Here, we characterize the effector of the Hippo signalling cascade, *yorkie*, during planarian regeneration and its role in restricting early injury responses. In *yki(RNAi)* regenerating animals, wound responses are hyper-activated; the bimodal proliferation kinetics are heighted and prolonged, while the transcriptional injury responses are similarly augmented with dysregulated temporal patterns. We also uncovered novel wound-induced genes by RNAseq that are primarily associated with tissue patterning. Indeed, a high proportion of non-wound- and wound-induced patterning molecules are mis-expressed in *yki(RNAi)*, which we demonstrate is in part due to an expanded muscle cell population. These altered injury responses have consequential effects on regenerative outcomes, specifically sensing the size of a given injury and appropriately scaling organ and tissue sizes. Taken together, our results suggest that *yki* functions as a key node to integrate the injury responses of proliferation, apoptosis, injury-induced transcription, and patterning to coordinate regeneration.

## Introduction

Regeneration is a growth-controlled program that is observed across the animal kingdom. Neo-natal mice can regrow missing digit tips, salamanders can replace missing limbs, while other organisms, such as hydrazoans and planarians, can regenerate virtually any tissue (Holstein et al., 2003; Rinkevich et al., 2011; Sanchez Alvarado, 2000; Takeo et al., 2013; Thornton, 1968). The shared characteristic to regenerate suggests a conserved genetic program that first replaces the missing tissue by proliferation (epimorphosis) and subsequently re-patterns and re-scales (morphallaxis) the regenerated tissue (Morgan, 1898; Morgan, 1901). Failure to initiate or cease aspects of either process can result in under- or overgrown tissues, respectively. Yet how proliferation and patterning are regulated in the correct spatiotemporal manner to determine size and scaling of regenerating tissues remains poorly understood.

The asexual planarian, *Schmidtea mediterranea*, displays a remarkable ability control growth because it can regenerate any missing tissue upon amputation, then rescale the entire animal in proportion to the remaining tissue. Planarian regenerative capacity is derived from its near ubiquitous population of adult stem cells (neoblasts), which are also the only mitotic cells in the animal and at least some of which are pluripotent (Reddien and Alvarado, 2004; Saló and Baguna, 1989; Wagner et al., 2011). In response to tissue removal, a stereotypical bimodal pattern of proliferation occurs first at 6 hours post amputation (hpa) and then at 48 hpa (Wenemoser and Reddien, 2010). Moreover, a common transcriptional injury response program is expressed with distinct temporal and spatial patterns (Wenemoser et al., 2012; Wurtzel et al., 2015). However, it is unknown how these two processes may be linked and what regulators may determine the onset or decay of these responses. As regeneration progresses, the missing tissues are replaced, reintegrated with the prexisiting tissue, and finally rescaled to restore tissue sizes and proportion. This rescaling process, also known as morphallaxis, is crucial to achieve proper proportions of the regenerated animal (Morgan, 1898; Morgan, 1901). For example, in planarians the WNT polarity gradients must be rescaled to accommodate the new size of the worm (Gurley et al., 2010). Therefore, the planarian is a unique model to understand the dynamic processes of growth—proliferation, patterning, and scaling—in an adult regenerative context.

The Hippo pathway is universal regulator of growth control and is a kinase cascade that impinges on the transcriptional co-activator, Yorkie, (in vertebrates: YAP1/2 and paralog TAZ) (Halder and Johnson, 2011; Huang et al., 2005). Constitutive activation of Yki or YAP results in overgrown tissues in flies and vertebrates, suggesting a conserved role in promoting cell division (Camargo et al., 2007; Dong et al., 2007; Huang et al., 2005). However, YAP can be growth-restrictive in highly proliferative and regenerative tissues, such as the mammalian intestine (Barry et al., 2012). Indeed, we have previously demonstrated that planarian *yorkie/YAP* (*Smed-yki*) is required to restrict stem cell proliferation, yet *yki(RNAi)* animals failed to regenerate (Lin and Pearson, 2014). This conundrum between increased proliferation with an undergrowth phenotype suggests other intricacies of growth control are dysregulated, but this remains to be tested. Here, we demonstrate that Yki is required for the regeneration of appropriately scaled and sized tissues by integrating the wound signals that direct the proliferative, apoptotic, and transcriptional injury responses. In *yki(RNAi*) animals, these known injury responses are hyper-activated with dysregulated patterns. Indeed, the proliferative dynamics are heighted and temporally prolonged. This enlarged stem cell population showed no block in differentiation, and surprisingly, showed increased numbers of both epidermal and *collagen*^+^ muscle cells. The expansion of these cell types can be attributed to a subsequent heighted transcriptional injury response. Using an RNA-deep sequencing (RNAseq) time course, we determined that *yki(RNAi)* regenerates had significantly dysregulated wound-induced genes, which have predominant roles in patterning. Thus, with altered proliferation kinetics and dysregulated expression of patterning molecules, *yki(RNAi)* animals failed to maintain proper size and scaling during morphallaxis. Altogether, this study demonstrates that the injury responses of proliferation and transcription are required to set up proper regenerative outcomes, such that *yki* is the crucial node that integrates these processes.

## Methods

### Animal husbandry, Exposure to γ-Irradiation, and RNAi

Asexual *Schmidtea mediterranea* CIW4 strain were reared as previously described (Sánchez Alvarado, 2002). For irradiation experiments, planarians were exposed to 60 Gray (Gy) of γ-irradiation from a ^137^Cs source (Pearson and Sánchez Alvarado, 2010). RNAi experiments were performed using previously described expression constructs and HT115 bacteria (Newmark et al., 2003). Bacteria were grown to an O.D.600 of 0.8 and induced with 1 mM IPTG for 2 hours. Bacteria were pelleted, mixed with liver paste at a ratio of 500 μ1 of liver to 100 ml of original culture volume, and frozen as aliquots. The negative control, “*control*(*RNAi*)”, was the *gfp* sequence as previously described (Cowles et al., 2013). All RNAi food was fed to one week starved experimental worms every 3^rd^ day for a total of 3 feedings. Amputations were performed 15 days after the final feeding with corresponding salt addition to suppress the homeostatic edema phenotype (Lin and Pearson, 2014). All animals were size-matched between experimental and control worms.

### Pharynx Amputations

Pharynxes were chemically amputated by replacing planarian water with 100 mM sodium azide (NaN_3_) diluted in planarian water. After 7-10 minutes of gentle swirling and pipetting, pharynxes were either entirely extruded or extended, which required forceps for complete removal. NaN3 was vigorously washed out with planarian media and subsequently fixed (Adler et al., 2014; Pellettieri et al., 2010).

### CHX experiments

Cycloheximide (Sigma) was added immediately to animals following amputation as previously described (Wenemoser et al., 2012).

### Quantitative Real-Time PCR (qRT-PCR)

Reverse transcription reactions were conducted on total RNA extracted from approximately 15 worms using a SuperScript III Reverse Transcriptase Kit (Invitrogen). Quantitative real-time PCR was performed in biological triplicate on a Bio-Rad CFX96 Touch^™^ Real-Tie PCR Detection System with SYBR Green PCR Master Mix (Roche) as per manufacturer’s instructions. The 2^−∆∆CT^ method was used for relative quantification. Primer pairs for ubiquitously expressed GAPDH were used as a reference as previously described (Eisenhoffer et al., 2008a) and regeneration time courses experiments were normalized to *control(RNAi)* intact levels.

### Immunolabeling, BrdU, TUNEL, and in situ hybridizations (ISH)

Whole-mount ISH (WISH), and double fluorescent ISH (dFISH), and immunostainings were performed as previously described (Currie et al., 2016a; Lauter et al., 2011; Pearson et al., 2009). Colorimetric WISH stains were imaged on a Leica M165 fluorescent dissecting microscope. dFISH and fluorescent phospho-histone H3 (rabbit monoclonal to H3ser10p, 1:500, Millipore) immunostains were imaged on a Leica DMIRE2 inverted fluorescence microscope with a Hamamatsu Back-Thinned EM-CCD camera and spinning disc confocal scan head. BrdU (Sigma B5002-5G, 25 mg/ml) was dissolved in 50% ethanol and fed to animals and was stained as previously described (Zhu et al., 2015). TUNEL was performed as previous described (Pellettieri et al., 2010) with the Terminal Deoxynucleotidyl Transferase enzyme (Thermo, EP0162). Cell counts and co-localizations were quantified using freely available ImageJ software (http://rsb.info.nih.gov/ij/). Significance was determined by a 2-tailed unequal variance pairwise student’s *t*-test. All images were post-processed in a similar manner using Adobe Photoshop.

### RNA sequencing and Differential Expression Analysis

RNA deep sequencing (RNAseq) was performed on 1 day post-irradiated animals with no amputation (intact) or tail fragments regenerating at 6, 12, and 24 hours post amputation. Experiments were performed in biological-triplicate, sequenced to a depth of >20 million reads per sample, and multiplexed on an Illumina HiSeq2500 with 50 base pair, single-end reads. Raw scRNAseq data from uninjured cells (including stem cells, neurons, gut, epithelial, muscle and peripharyngeal cells) were obtained from the NCBI Sequence Read Archive (SRA:PRJNA276084) (Wurtzel et al., 2015). Reads were aligned to the SmedASXL transcriptome assembly under NCBI BioProject PRJNA215411 using bowtie2 (Langmead and Salzberg, 2012) with 15 bp 3’ trimming. For validating and detecting novel wound-induced genes, DEseq2 was used for differential expression analysis on triplicated regenerating samples at each time point compared to intact animals (Love et al., 2014). The parameters for significance calling were [FC] (fold change) > 1.2 with a [FDR] (false discovery rate) < 0.05. To detect mis-expressed patterning molecules, DEseq2 was run on the triplicated *yki(RNAi)* samples and the matched *control(RNAi)* with [FC] > 1.2 or [FC] < 0.83 with a [FDR] < 0.05. Violin plots were produced using modified source code from (Macosko et al., 2015) and heatmaps were produced using the modified heatmap.3 source code from (Molinaro and Pearson, 2016).

## Results

### yorkie regulates stem cell proliferation during regeneration

We previously reported that *yki* is required for planarian regeneration, but the underlying cause remained to be elucidated. Over 90% of *yki(RNAi)* tail fragments showed no evidence of a blastema at 5 days post-amputation (dpa) and failed to regenerate, while head and trunk fragments failed to a lesser degree (Lin and Pearson, 2014). Because tail fragments were the most severely affected, we chose to focus our characterization on tails.

Failures in regeneration are often correlated with aberrant stem cell dynamics, including stem cell proliferation. Thus, we first examined proliferation with mitotic marker phosphorylated histone 3 (H3P). In a *control(RNAi)* regeneration time course, proliferation occurred in a bimodal pattern with two bursts of proliferation that occurred at 6 and 48 hours post amputation (hpa), as previously described (Figure 1A) (Wenemoser and Reddien, 2010). However, in *yki(RNAi)* animals, this bimodal pattern was perturbed temporally and in its amplitude. The first wave of proliferation was prolonged and peaked 15 hours later at 24 hpa. The onset of second wave of proliferation was delayed and peaked at 72 hpa in *yki(RNAi)* animals (Figure 1B). Therefore, *yki* was required to restrict the proliferation response to injury, which was also observed in head and trunks (Supplemental Figure 1A-D).

**Figure 1.**
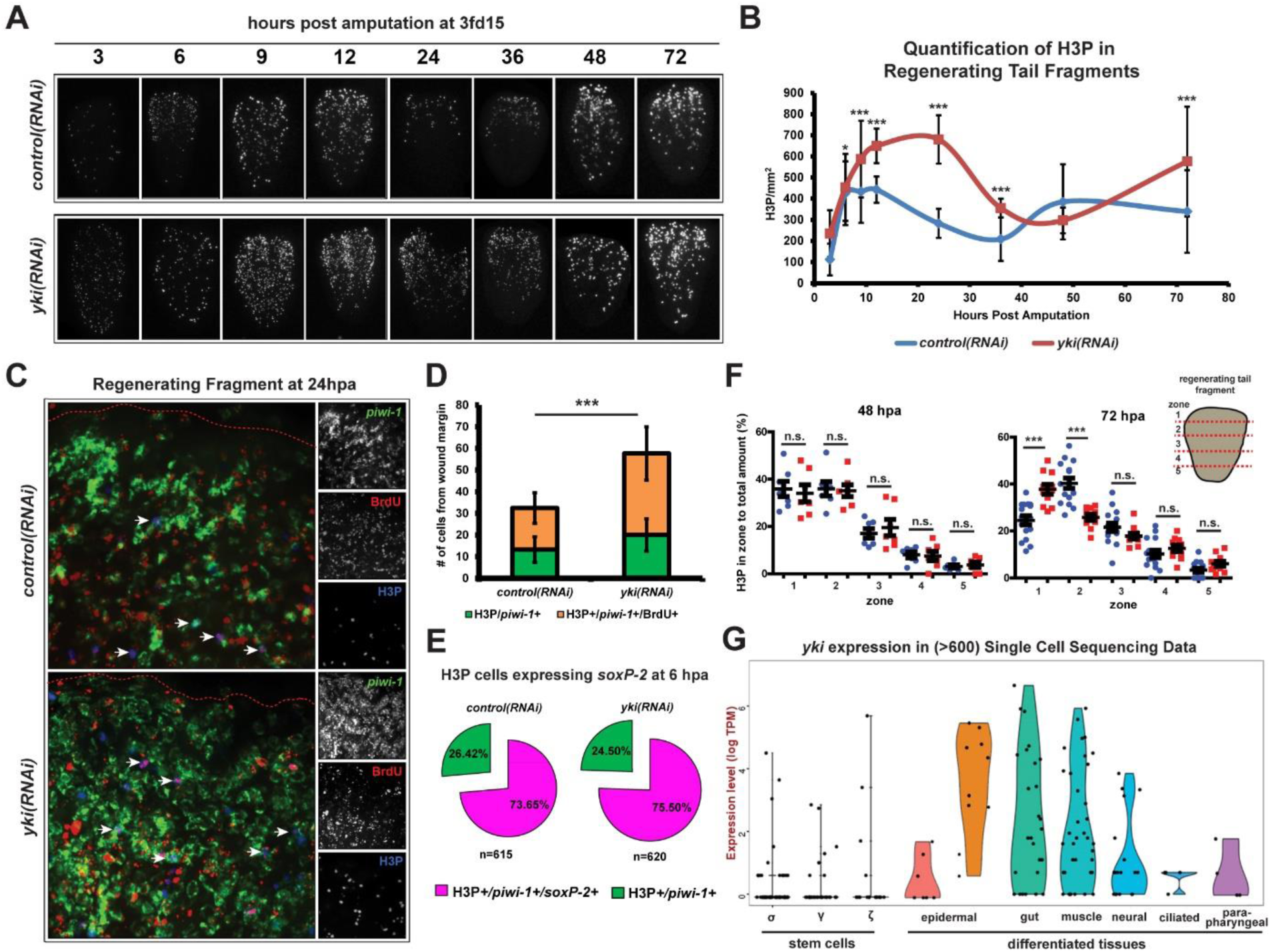
*yki* regulates stem cell proliferation dynamics during regeneration. (A) Time course of regenerating fragments stained with mitotic marker, phosphorylated histone 3 (H3P). (B) The quantification of time course in A. (C) *yki(RNAi)* does not affect the cell cycle. Triple positive cells (H3P+/*piwi-1*+/BrdU+) are marked with white arrows. Red dotted line is the wound boundary. (D) No significant difference is observed (*p*=0.67) in the percentage of H3P+/*piwi-1*+/BrdU+ cells (orange bar) to the total amount of H3P cells (orange and green bars summed) in *control(RNAi)* compared to *yki(RNAi)* tail fragments (59±9% and 60±7%, respectively). (E) Percentages of H3P+/*piwi-1*+ cells expression sigma stem cell marker *soxP-2* (pink) or not (green). (F) The percentage of mitoses occurring in each zone compared to the total amount at 48 and 72 hpa in *control(RNAi)* (blue) and *yki(RNAi)* (red). Zones are divided into 5 equal compartments (20% each) with zone 1 at the anterior, closest to the wound site. (G) From single-cell RNAseq, *yki* expression is enriched in the differentiated tissues (epidermal, gut, and muscle), and is lowly expressed in the stem cell compartment. Data show mean ± s.d. n.s.=not significat, \**p*<0.05, **<*p*<0.01, \*\*\**p*<0.001.

Canonical target genes for *Drosophila* Yki are cell cycle regulator *cyclin E* and apoptosis inhibitor *diap1* (Huang et al., 2005). Thus, the increased proliferation in *yki(RNAi)* could be attributed to a change in the cell cycle. Using BrdU pulse-time-chase experiments, we examined the proportion of label retaining cells at the wound margin (Figure 1C). The quantification of S-phase (BrdU^+^) or G2/M-phase (H3P^+^) cells suggested that *yki* does not inhibit the entry or exit into S-phase. Moreover, the ratio of label retaining cells (H3P^+^/*piwi-1*^+^/BrdU^+^) to the total amount of H3P cells was not different between *control(RNAi)* and *yki(RNAi)*, 59±9% and 60±7%, respectively, despite higher proliferation in *yki(RNAi)* (Figure 1D). If *yki* affected the cell cycle, a difference in the percentages would be expected, however, the opposite was seen. Therefore, the changes in proliferation in *yki(RNAi)* animals cannot be attributed to alterations in the length of the S or G2 phase individually, although we cannot rule out the possibility that the whole cell cycle is accelerated. Finally, the increased proliferation did not change stem cell subclass identities (Figure 1E) (van Wolfswinkel et al., 2014) and proliferating cells were also not undergoing apoptosis because no H3P^+^/TUNEL^+^ cells were observed (Supplemental Figure 1E-F).

In response to amputation, a wave-like pattern of proliferation emanates from the wound site over time, suggesting a diffusion-based mechanism that triggers proliferation (Elliott and Alvarado, 2013; Wenemoser and Reddien, 2010). For instance, proliferation predominantly occurs at the wound margin at 48 hpa and subsequently shifts posteriorly by 72 hpa (Figure 1F). However, in *yki(RNAi)*, proliferation was sustained at the wound margin at 72 hpa (Figure 1F). This suggested the possibility of a cell non-autonomous proliferative cue that is upregulated in *yki(RNAi).* Indeed, we have previously reported that *yki* was not enriched in the stem or progenitor cell populations by bulk RNA sequencing (Lin and Pearson, 2014). Congruent with our previous results, single cell RNA sequencing (scRNAseq) of stem and differentiated tissues also demonstrated that *yki* was not enriched in any of the stem cell sub-classes (γ, ζ or σ), and instead, is primarily expressed in differentiated tissues, such as the epidermis, muscle, and gut (Figure 1G). Interestingly, these same differentiated tissues have recently been shown to be enriched for the expression of wound-induced genes, collectively known as the transcriptional injury response (Wurtzel et al., 2015). Therefore, we next tested whether defects in the early transcriptional response to wounds could explain defects observed in *yki(RNAi)* regenerates.

### *yki* is required to restrict the transcriptional injury response

In planarians, the transcriptional wounding response program is activated immediately following any type of injury, whether tissue is removed—such as a wedge cut and an amputation—or not, like an incision and a poke (Wenemoser et al., 2012). This response can be subcategorized temporally into two waves of transcription, an early immediate wave that peaks in expression by 6 hpa, and a subsequent late sustained wave that prolongs in expression until 24 hpa (Wenemoser et al., 2012; Wurtzel et al., 2015). In *yki(RNA)*, *fos-1*, an early wave marker, was precociously expressed earlier at 0.5 hpa and was prolonged until 12 hpa (Figure 2A). Likewise, the late marker *delta-1* was also increased and temporally unrestricted in its expression (Figure 2B). The time courses observed by WISH were supported by parallel experiments using qRT-PCR (Figure 2C–D). Two other wound-induced genes, *jun-1* and *tyrosine-kinase2*, showed similar, upregulated expression in regenerating *yki(RNAi)* head, trunk, and tail fragments (Supplemental Figure 2A-B). Additionally, irradiated animals showed no difference in *fos-1* or *delta-1* expression compared to their non-irradiated controls, suggesting that the increased mitoses in *yki(RNAi)* did not contribute to the elevated wound-induced gene expression (Figure 2E-G). Furthermore, *yki* itself was not injury-induced (Supplemental Figure 2C) and cycloheximde, which blocks protein translation, did not alter the transcriptional response (Supplemental Figure 2D-E) (Wenemoser et al., 2012). Finally, edemas did not affect the proliferative or transcriptional injury responses, demonstrating that the pleiotropic effects of *yki* on the excretory system were independent (Supplemental Figure 3) (Lin and Pearson, 2014). Due to the specificity of dysregulated wound-response transcripts in *yki(RNAi)* fragments, we next tested whether this could be used to discover novel wound-induced programs.

**Figure 2.**
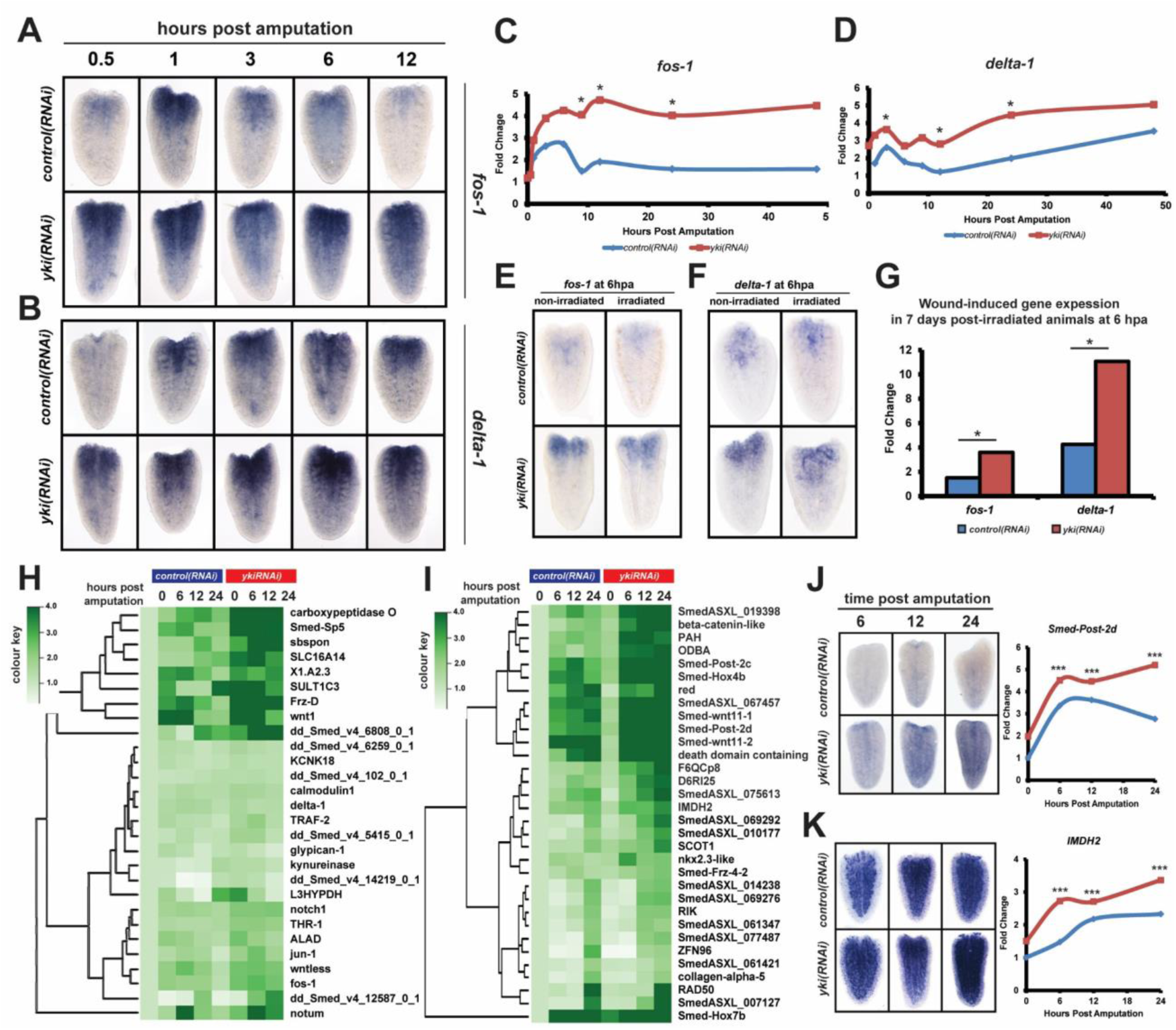
*yki* is required to restrict the transcriptional injury response. (A-B) A WISH time course of wound-induced genes *fos-1* (A) and *delta-1* (B) during regeneration. (C-D) qRT-PCR experiments parallel results by WISH in A. (E-G) Irradiated worms show the same *fos-1* (E) and *delta-1* (F) response as non-irradiated worms by WISH (E-F) and qRT-PCR (G). (H) A heatmap of significantly upregulated (fold change [FC] > 1.2, false discovery rate [FDR] < 0.05, pair-wise comparison between matched time points) *bona fide* wound-induced genes normalized to *control(RNAi)* intact levels (I) Heatmap of novel wound-induced genes with [FC] normalized to *control(RNAi)* intacts. (J-K) Regeneration time course stained by WISH (left) with corresponding CPM fold-change normalized to *control(RNAi)* intact (right) for: *Smed-Post-2d* (J) and *IMDH2* (K). \**p*<0.05, \*\**p*<0.01.

### The identification of novel wound-induced genes by RNAseq

The loss of restriction in the transcriptional injury response in *yki(RNAi)* suggested that transcriptome profiling *yki(RNAi)* animals may uncover novel wound-induced genes. Previous studies have focused on isolating and sequencing small wound pieces instead of the whole amputated fragment. Although this approach has been very successful (Wenemoser et al., 2012; Wurtzel et al., 2015), it would removes candidates whose expression may be induced away from the wound site. Moreover, we chose to irradiate animals because *yki(RNAi)* caused an expanded stem cell population, which could greatly alter many non-injury response transcripts (Figure 1A–B) (Lin and Pearson, 2014). Therefore, we conducted RNA-deep sequencing (RNAseq) on 1 day post-irradiated intact animals as well as regenerating tails at 6, 12, and 24 hpa.

First, we conducted validation on a set of 128 previously identified wound-induced genes (Wurtzel et al., 2015). We curated this list to 97 transcripts for our analyses, removing stem cell enriched transcripts because our samples were irradiated. By pair-wise DEseq2 comparison of *yki(RNAi)* to the respective *control(RNAi)* time point, we identified 32 of the 97(33%) transcripts significantly upregulated (fold change [FC] > 1.2, false discovery rate [FDR] < 0.05) at any time point (Supplemental Table 1). By normalizing the expression to *control(RNAi)* intacts, an elevated response was evidently observed in *yki(RNAi)* tail fragments (Figure 2H). Consistent with the above WISH and qRT-PCR data, *jun-1*, *delta-1*, and *fos-1* were also significantly upregulated by these analyses (Figure 2H). We further validated the upregulation in *yki(RNAi)* with another *bona fide* injury marker that we had not previously examined, *Smed-sp5*, which is a downstream target of *Smed-ß-catenin* that is also wound-induced (Reuter et al., 2014; Wurtzel et al., 2015). We observed that *Smed-sp5* was precociously expressed in *yki(RNAi)* animals at 6 hpa and was sustained well beyond its normal temporal peak (Supplemental Figure 4A).

In order to find novel wound-induced genes, we used DEseq2 to compare RNAseq from *control(RNAi)* intact worms to 6, 12, and 24 hpa tail fragments. This analysis generated a list of 341 significantly upregulated transcripts ([FC] > 2, FDR < 0.05). Using this list, we compared *yki(RNAi)* to *control(RNAi)* regenerates at their matched time points ([FC] > 1.25, FDR <0.05), which resulted in a list of 42 transcripts, only 9 of which have been previously characterized. Interestingly, multiple patterning molecules were identified, including HOX proteins *Smed-Post-2c*, –*2d*, *Hox4b*, *Hox7c*, and WNT-signalling pathway genes *wnt-11-1*, -*11-2*, and *Frz-4-2* (Figure 2I) (Supplemental Table 2). *Smed-Post-2d*, which is similar to an AbdominalB-like homeobox protein, was wound-induced with expression that peaked at 12 hpa (Figure 2J) (Currie et al., 2016a; Iglesias et al., 2008). By contrast, in *yki(RNAi)*, the response was induced earlier, heighted, and sustained (Figure 2J). Moreover, this trend was also observed for *IMDH2* (Figure 2K) and *beta-catenin-like* (Supplemental Figure 4B). Interestingly, by scRNAseq analyses, all 3 novel candidates had expression in the muscle, while *Smed-Post-2d* and IMDH2 also had predominant enrichment in the epidermal (early and late) progenitor populations (Supplemental Figure 4C) (Wurtzel et al., 2015). Therefore, we next examined the epidermal and muscle cell populations for defects in *yki(RNAi)* animals due to multiple dysregulated wound-induced genes were expressed in these tissue types (Supplemental Figure 4C-D).

### *yki(RNAi)* animals show higher epidermal density and smaller cell size

The epidermis is enriched for wound-induced genes and has a well-defined lineage that can be readily assayed (Eisenhoffer et al., 2008b; Tu et al., 2015; van Wolfswinkel et al., 2014; Wenemoser et al., 2012; Wurtzel et al., 2015; Zhu et al., 2015). Early epidermal progenitors marked by *prog-2* were significantly increased in density in *yki(RNAi)* at 2 dpa. Similarly at 7 dpa, *yki(RNAi)* showed more *AGAT-1*^+^ late epidermal progenitors and differentiated dorsal epidermal cells compared to *control(RNAi)* (Figure 3A-B). This suggested that epidermal differentiation is not blocked in *yki(RNAi)*, and instead, *yki* was required to restrict epidermal numbers. The increased epidermal density in *yki(RNAi)* came at the expense of the cell size (Figure 3C) and accordingly, the cell cortices of the epidermis were also smaller by Concanavalin A staining (Figure 3D) (Zayas et al., 2010). This suggested that the increased epidermal density may contribute to elevated wound-induced gene expression.

**Figure 3.**
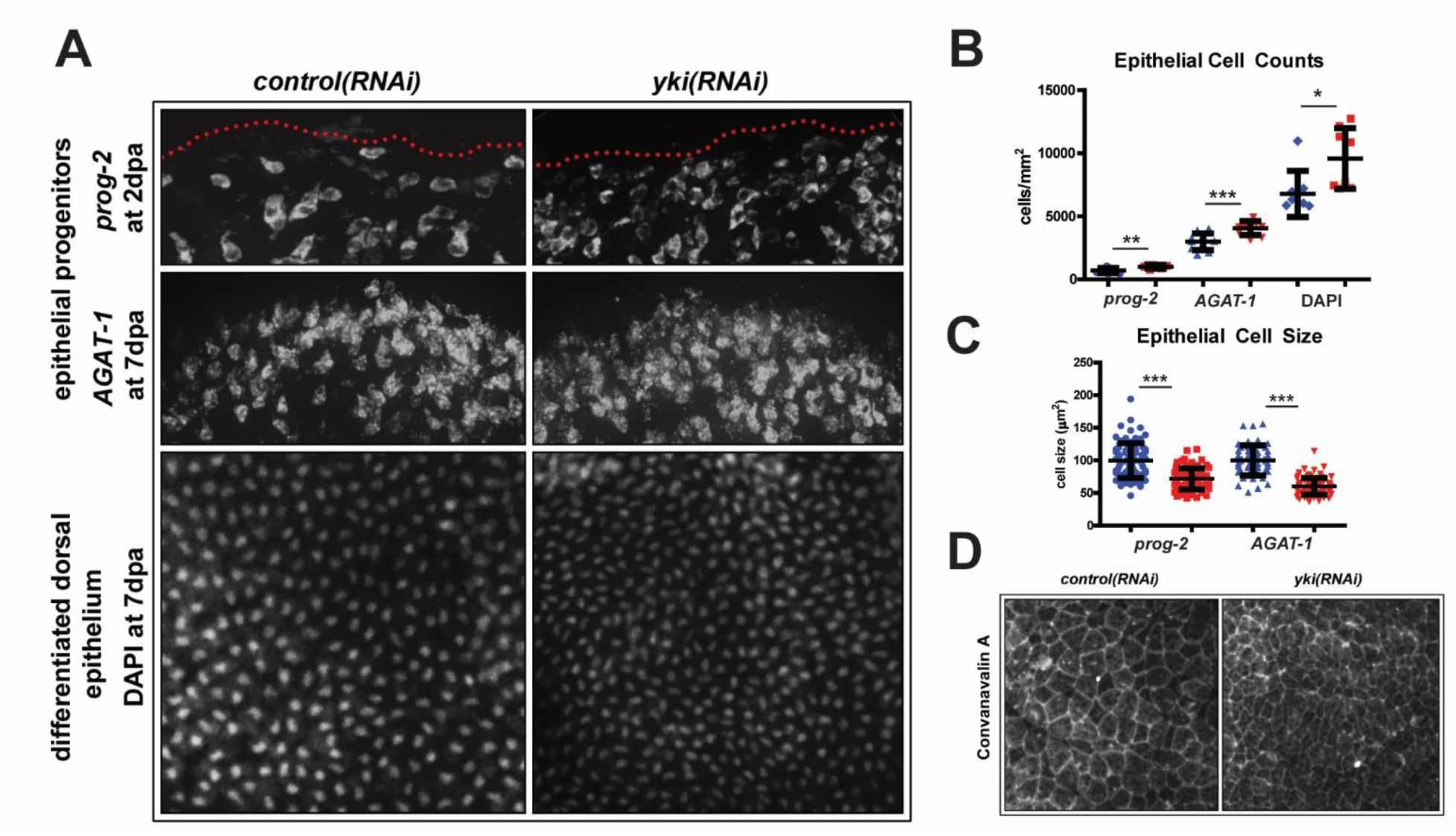
*yki* restricts epidermal density and cell size. (A-C) Epidermal populations assayed with *prog-2* at 2 dpa, and *AGAT-1* and DAPI at 7 dpa by WISH (A). Red dotted line indicates wound margin. Histogram of epidermal cell counts (B) and cell sizes (C) by quantifying images in A. Blue symbols are *control(RNAi)* and red is for *yki(RNAi).* (D) Epidermal junctions stained with Concanavalin A. Data show mean ± s.d. \**p*<0.05, \*\**p*<0.01, \*\*\**p*<0.001.

### An expanded population of muscle cells and patterning molecules are observed in yki(RNAi) animals

We next focused on the muscle population because it was the tissue with the most dysregulated wound-induced genes in *yki(RNAi)*. Moreover, the *collagen*^+^ muscle subpopulation is the source of many patterning signals that predominantly belong to major signalling cascades such as WNT, BMP, and TGF-β (Scimone et al., 2016; Witchley et al., 2013). In order to understand the dynamics of the muscle cell population during regeneration, we administered BrdU at 3fd15, amputated worms 5 days afterwards, and subsequently fixed them at 7 dpa (Figure 4A). In *yki(RNAi)* animals, an increased number of *collagen^+^/BrdU^+^* cells was observed compared to controls (Figure 4B-C). Accordingly, we found that *collagen* and *troponin*, another muscle-specific gene, were upregulated immediately after injury in *yki(RNAi)* tail fragments by qRT-PCR (Figure 4D). Thus, the increased muscle cell population could also account for the augmented and aberrant transcriptional wound response in *yki(RNAi)*.

**Figure 4.**
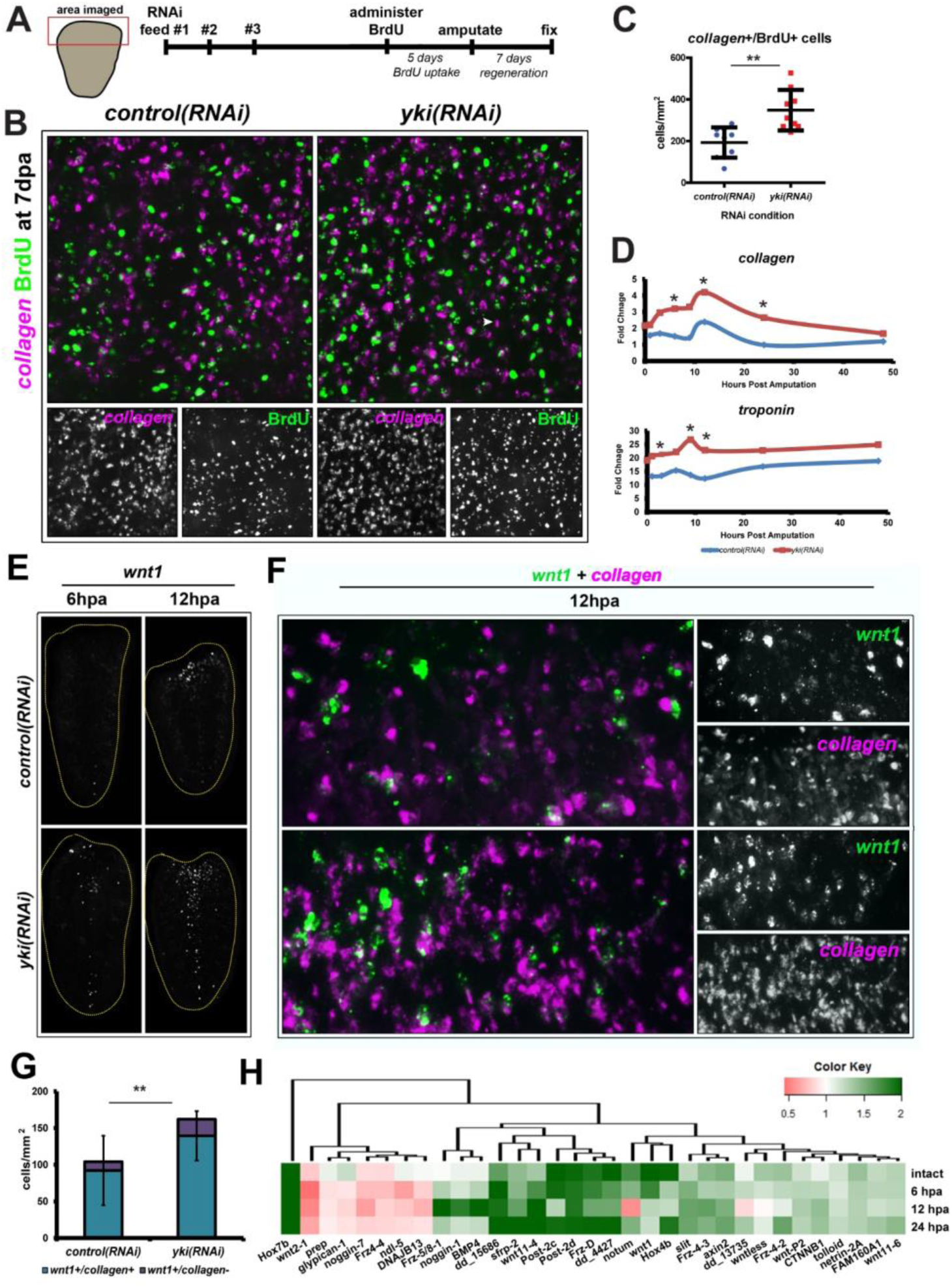
*yki(RNAi)* causes an expansion of the muscle population and mis-expression of patterning molecules, including *wnt-1.* (A-C) BrdU chases into the *collagen+* muscle population. A timeline of the BrdU experiment (A). FISH for *collagen* (magenta) and BrdU (green) (B) with double positive cells quantified in a histogram (C). (D) qRT-PCR for *collagen* and *troponin* during a regeneration time course. (E) *wnt1* expression at 6 hpa and 12 hpa by WISH. Yellow dotted line outlines the worm fragment. (F) dFISH of *wnt1* (green) and *collagen* (magenta). (G) A quantification of *wnt1+* cells that are *collagen+* (turquoise) and *collagen-* (purple) at 12 hpa. A significant difference in the total number (turquoise + purple bars summed) is observed in *yki(RNAi).* (H) The fold change of significantly changing patterning genes in *yki(RNAi)* compared to *control(RNAi)* in intacts, 6, 12, and 24 hpa (FDR < 0.05). Data show mean ± s.d. *p<0.05, **p<0.01.

To test whether the overproduction of muscle cells in *yki(RNAi)* was attributed to increased wound-induced gene expression, we examined *wnt1*. Injury induces *wnt1* expression, but it is also a polarity determinant that has >90% co-localization within the *collagen*+ muscle cell population (Petersen and Reddien, 2009; Witchley et al., 2013). In *yki(RNAi)* animals at 6 and 12 hpa, significantly more *wnt1*^+^ cells were observed along the tail midline, but also ectopically at the wound margin (Figure 4E). These ectopic *wnt1*^+^ cells were predominantly *collagen*^+^ with no significant difference in the percentage of *wnt1*^+^/*collagen*^−^ cells in *yki(RNAi)* as compared to *control(RNAi)* (14.2±3.9% and 11.8±8.1%, respectively; *p*=0.39) (Figure 4F-G). Therefore, the increased *wnt1*^+^ cells in *yki(RNAi)* tails were not ectopically expressed in a different cell type and suggested that the expanded muscle population can be attributed to the expanded expression of wound-induced genes.

Many non-wound-induced body patterning molecules are also known to be highly expressed in *collagen*^+^ cells, including the HOXs and FGFs (Scimone et al., 2016; Witchley et al., 2013). With the increased muscle population in *yki(RNAi)* animals, we tested whether other wound and non-wound induced patterning molecules were aberrantly expressed. By pair-wise DEseq2 analyses between *yki(RNAi)* and *control(RNAi)* matched time points, we found 35 patterning genes significantly changing at any given time point ([FC] > 1.2 or [FC] < 0.83 with [FDR] < 0.05) (Figure 4H and Supplemental Table 3). A high proportion of these genes were associated with WNT signalling—a key determinant in anterior-posterior identity—which was expected because *yki(RNAi)* tails do not regenerate their anterior (Lin and Pearson, 2014). However, *notum*, which antagonizes WNT signalling to promote head specification, as well as the anteriorly-expressed *sfrp-2* were still expressed in *yki(RNAi)* tails (Figure 4H and Supplemental Figure 5A-B). Thus, the *yki(RNAi)* regenerative defects were not simply a failure to express anterior-specification genes. Indeed, the changes in patterning were not limited to the WNTs, but also included molecules associated with patterning the midline such as *slit-1* and *netrin-2A* (Supplemental Figure 5C), or the dorsal-ventral axis such as *bmp, tolloid-1, noggin-1*, and *-7* (Figure 4H). Alterations in expression of signalling pathways can affect tissue maintenance with improperly sized organs, thus, we next tested whether *yki(RNAi)* animals had defects in organ scaling (Hill and Petersen, 2015; Reddien, 2011; Scimone et al., 2016).

### *yki(RNAi)* animals exhibit defects in scaling and morphallaxis

The planarian has an innate ability to sense injury size by recognizing the amount of tissue removed, known as the “missing tissue response” or “size sensing mechanism” (Gaviño et al., 2013). We wanted to test whether this mechanism was still in place in *yk(RNAi)* animals because the injury responses were heightened, prolonged, and sustained (Figures 1 and 2). The regenerating planarian pharynx is an ideal organ to test for the animal’s ability to sense the size of wounds. In relation to the amount of tissue removed or amputated, the post-mitotic planarian pharynx must be rescaled and re-patterned to its new fragment size, which is accomplished in part by changes in cell death (Pellettieri et al., 2010). To test how this scaling mechanism may be altered in *yki(RNAi)*, pharynxes were chemically amputated from worms that had been regenerating for 3 days but underwent selective amounts of tissue removal (0%, 10%, 40%, and 80%) (Figure 5A). The isolated pharynxes were immediately fixed and assayed for TUNEL (Figure 5B). In *control(RNAi)* amputated pharynxes, a proportionality between the amount of tissue removed and the amount of cell death was observed, as previously reported (Figure 5B). However, in *yki(RNAi)*, this relationship was uncoupled (Figure 5B-C). This suggested that *yki* was required to mediate a scaling process or injury-size sensing mechanism during the initial stages of regeneration.

**Figure 5.**
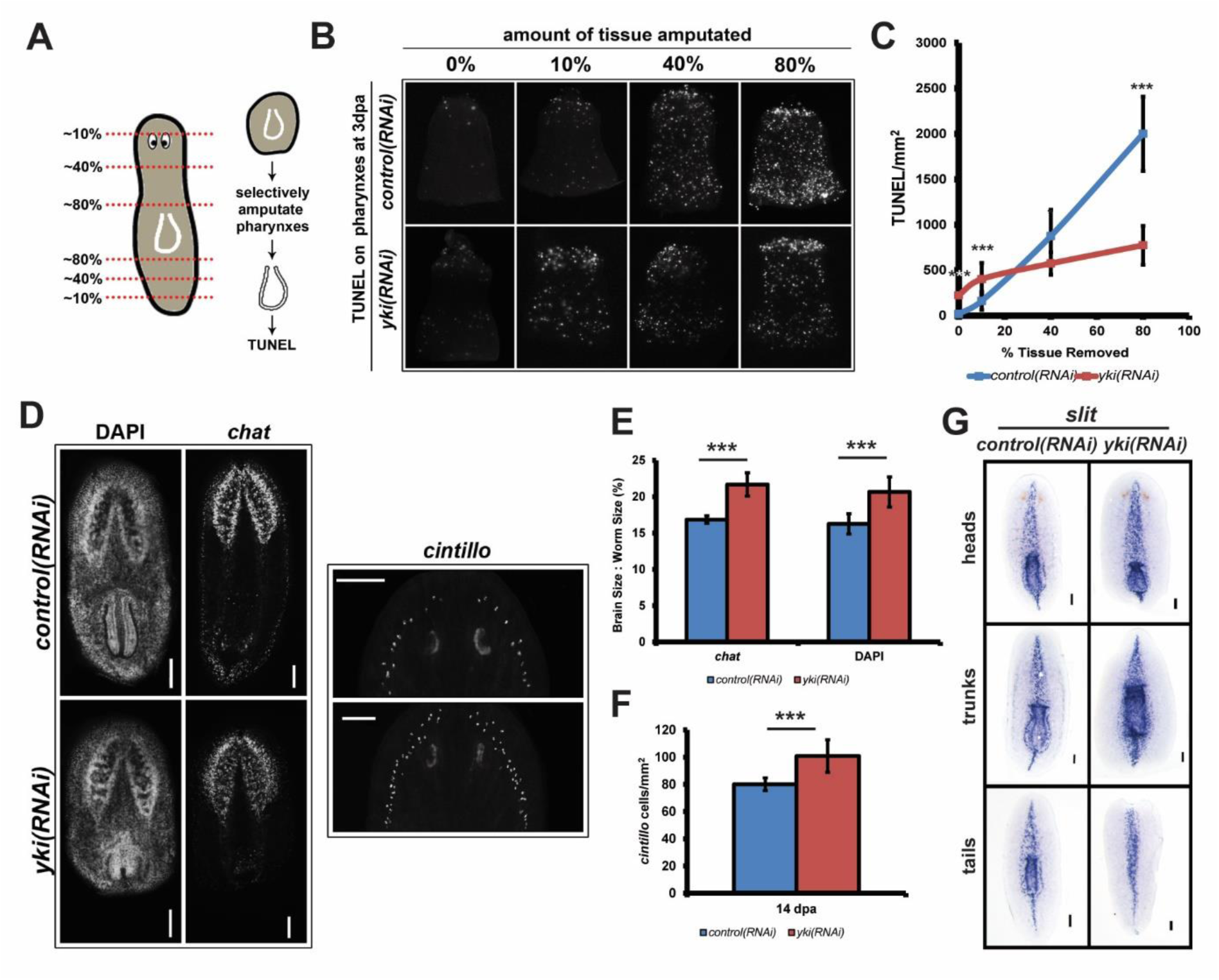
*yki* regulates scaling and morphallaxis. (A) Pharynxes amputated from animals with selective tissue removed can be assayed for cell death as a size sensing mechanism. A schematic of the regeneration series with various percentages of tissue removed with subsequent experimental steps outlined. (B) Amputated pharynxes assayed for TUNEL. (C) Quantification of TUNEL stained pharynxes from A. (D) Head fragments from *control(RNAi)* and *yki(RNAi)* worms were assayed for brain size and morphology with DAPI, *chat,* and *cintillo* at 14 dpa. Scale bar is 100 μm. (E) The ratio of brain size to worm size expressed as a percentage and quantified from D. (F) Quantification of the number of *cintillo+* cells to worm size from E. (G) Regenerating fragments at 14 dpa stained with *slit. yki(RNAi)* animals show an expanded *slit* gradient that fails to rescale to appropriate proportions. Data show mean ± s.d. \**p*<0.05, \*\*\**p*<0.001.

The role for *yki* to mediate scaling can also be assayed during the process of morphallaxis or re-scaling of body and tissue proportions. For example, similar to the pharynx in trunk fragments, the planarian brain must be scaled down in size in relation to the new size of the head fragment (Hill and Petersen, 2015; Morgan, 1898; Reddien and Alvarado, 2004). Using DAPI and *chat (choline acetyltransferase)*, which specifically labels mature cholinergic neurons, a ratio of the brain size relative to the body size was determined (Figure 5D) (Agata et al., 1998). At 14 dpa, *yki(RNAi)* head fragments were smaller due to a regeneration defect (Lin and Pearson, 2014), however, its brain to body size was significantly larger (Figure 5E). Accordingly, *cintillo,* which is expressed in chemoreceptive neurons, also did not maintain scale in *yki(RNAi)* animals (Figure 5F) (Hill and Petersen, 2015; Sanchez Alvarado et al., 2002). Similarly, the *slit* midline gradient was also enlarged in *yki(RNAi)* at 14 dpa in all three regenerating fragments (Figure 5G) (Cebria et al., 2007). Finally, *yki(RNAi)* animals with an eye injury showed a heightened transcriptional injury response and regenerated significantly larger compared to the contralateral uninjured one (Supplemental Figure 6). Taken together, this suggested that *yki* was required to determine organ scaling by restricting the magnitude and duration of injury responses (Figure 6).

**Figure 6.**
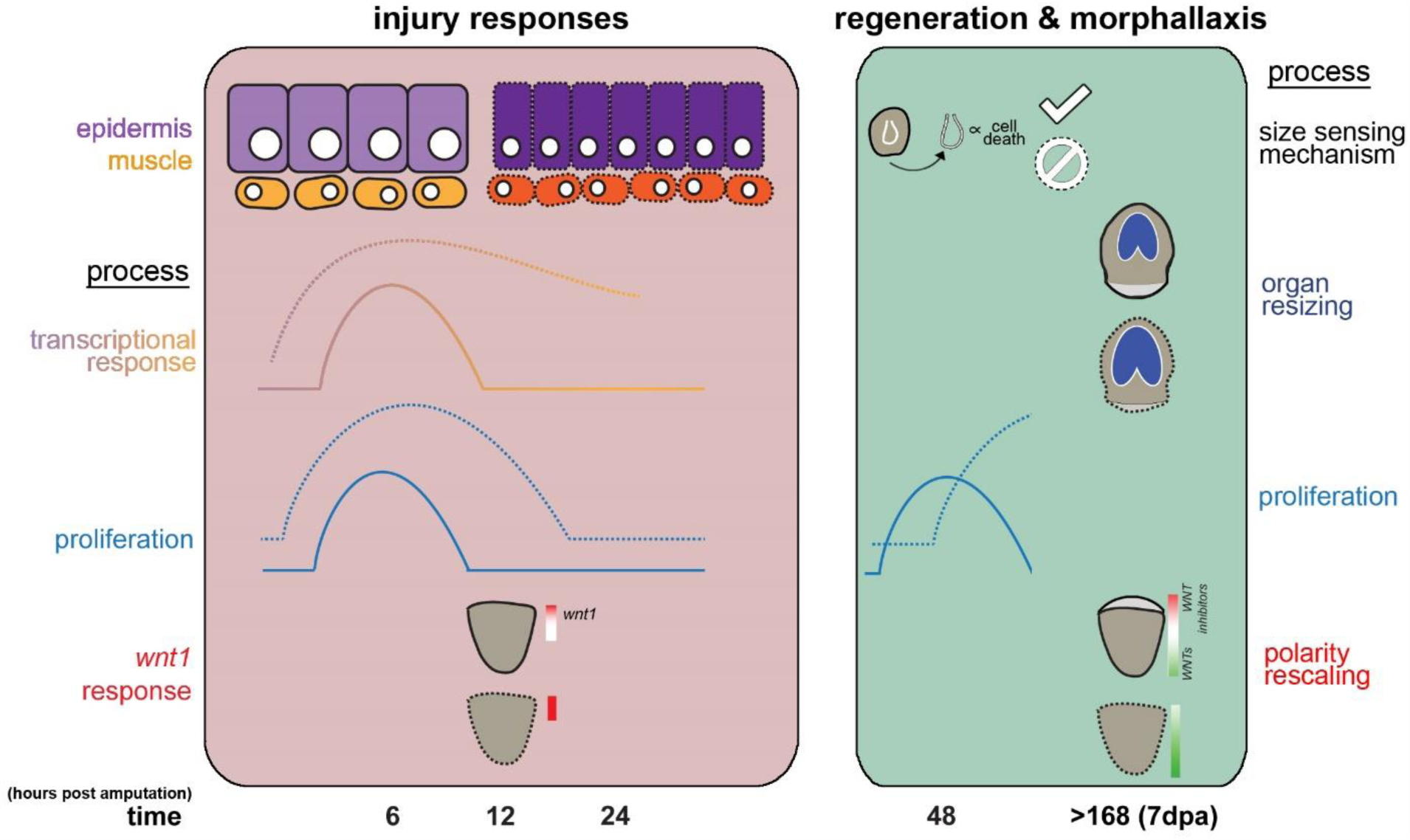
Figure 6. The roles for *yki* during planarian regeneration. In *control(RNAi)* animals (solid lines), the injury responses are composed of distinct spatial and temporal patterns of proliferation and transcription. For instance, the transcriptional response of wound-induced genes is primarily localized to the epidermis and muscle, while the *wnt1* response peaks at 12 hpa. As regeneration progresses, a second burst of proliferation occurs at 48 hpa. After the lost tissues have been replaced, reintegration with the old tissues occurs by morphallaxis. For example, the brain will be rescaled down in size in head fragments, and the WNT genes re-establish their graded expression along the anterior-posterior axis. By contrast, in *yki(RNAi)* regenerating animals (dotted lines), all aspects of the generic wound response are heightened, evident by dysregulated proliferative dynamics and the augmented wound-induced gene expression, likely attributed to the increased epithelial and muscle cell numbers. The second mitotic burst is temporally shifted and a failure in regeneration is observed. Ultimately, repatterning and scaling are also affected. Altogether, this suggests that *yki* is required to restrict multiple aspects the injury response, including proliferation, wound-induced gene expression, and patterning to ensure proper regenerative outcomes.

## Discussion

Regeneration is a complex, multistep process that requires precise spatial and temporal regulation. The first step is an injury response comprised of a burst of proliferation, apoptosis, and a transcriptional injury response, which together provide the initial basis for planarian regeneration (Pellettieri et al., 2010; Wenemoser et al., 2012; Wenemoser and Reddien, 2010; Wurtzel et al., 2015). The transcriptional injury response is a generic and robust program that is activated immediately following any type of injury, whereby failures of activation have ramifications during regeneration (Almuedo-Castillo et al., 2014; Wenemoser et al., 2012; Wurtzel et al., 2015). Here, we demonstrate that *yki* is required to restrict this response (Figure 6). In all three *yki(RNAi)* regenerating fragments, wound-induced genes were precociously expressed earlier with a heighted response evident by WISH, qRT-PCR, and RNAseq (Figure 2A-K). This aberrant signalling of mitogens, transcription factors, and patterning molecules may subsequently affect the proliferative kinetics. Although the signal(s) that induce the bimodal waves of proliferation remain elusive in planarians, correlations between the transcriptional and proliferative response may be inferred. First, the two responses overlap temporally (Wenemoser et al., 2012; Wenemoser and Reddien, 2010; Wurtzel et al., 2015). Second, *follistatin(RNAi)* regenerating animals fail to express a second wave of wound-induced genes and also fail to mount a second burst of proliferation, suggesting a possible link between the two processes (Gaviño et al., 2013; Roberts-Galbraith and Newmark, 2013). In addition, previous work profiling liver regeneration identified waves of transcription that prime hepatocytes for subsequent proliferation (Fausto, 2000; Haber et al., 1993; Su et al., 2002). Therefore, a similar mechanism may be occurring in planarians, whereby the transcriptional injury response is required to initiate proper regenerative proliferative kinetics.

Along with an altered transcriptional injury response, *yki(RNAi)* animals also showed dysregulated proliferation patterns. Normally, proliferation occurs in a bimodal wave (Wenemoser and Reddien, 2010), however in *yki(RNAi)*, the first peak was significantly increased and prolonged, while the second one was delayed and similarly heighted (Figure 1A-B). These altered temporal patterns are reminiscent to the changes also observed in the transcriptional injury response, suggesting a possible relationship between the two. Moreover, the changes in proliferation may not be cell autonomous because *yki* is not enriched the stem cells (Figure 1G) (Lin and Pearson, 2014; Wurtzel et al., 2015). The stem cell subclass identities of the proliferating population remained unchanged in *yki(RNAi)*, suggesting that *yki* does not affect stem cell lineage commitment. Indeed, differentiation proceeded at even higher levels in *yki(RNAi)* (Figure 3, epidermal; Figure 4, muscle). Therefore, the global increase in proliferating stem cells may be attributed to a cell non-autonomous factor, perhaps from the increased transcriptional injury response. Congruent with this hypothesis, proliferation was sustained at the wound margin at 72 hpa in *yki(RNAi)*, whereas it normally occurs more posteriorly in *control(RNAi)* at this time point (Figure 1E). Perhaps this role of *yki* in planarians is similar to *Drosophila* midgut regeneration where Yki is activated in the non-proliferative enterocytes to secrete cytokines that stimulate intestinal stem cell proliferation in a non-autonomous fashion (Karpowicz et al., 2010; Shaw et al., 2010; Staley and Irvine, 2010). Similarly, axolotl limb regenerative proliferation is regulated by non-autonomous cues, such as the secreted molecule MARCKS that is also transcriptionally injury-induced, although it is unknown whether a *yorkie* homolog is involved (Sugiura et al., 2016). Future studies elucidating what the injury-induced molecule that triggers proliferation would provide great insights into initiation of regenerative programs in planarians.

Another key feature of the transcriptional injury response is to lay the foundation for axial and tissue re-scaling. The 12 hpa time point is the peak temporal expression window for the majority of wound-induced patterning molecules (Wenemoser et al., 2012; Wurtzel et al., 2015), including *wnt1* which is crucial in determining axial polarity (Petersen and Reddien, 2009). Interestingly, many of the dysregulated wound-induced genes in *yki(RNAi)* have roles in patterning (Figure 2H-I). Indeed, *wnt1* was significantly increased in *yki(RNAi)* animals (Figure 3G). Modulating patterning can have consequences in regeneration by affecting pole determination, cell fate decisions, or organ sizing and scaling (Currie et al., 2016b; Gurley et al., 2008; Hill and Petersen, 2015; Lander and Petersen, 2016; Scimone et al., 2016; Scimone et al., 2014). We hypothesize that the failures in regeneration in *yki(RNAi)* may be attributed to the mis-expression of multiple patterning molecules (Figure 4H). In addition, the process of morphallaxis was also altered in *yki(RNAi)* (Figure 5D-G). Therefore, the dysregulated transcriptional injury response in *yki(RNAi)* may be responsible for the changes in proliferation and patterning that likely have detrimental effects on scaling and sizing during regeneration (Figure 6).

What could be causing the upregulation of the transcriptional injury response in *yki(RNAi)* animals? One possibility may be epithelial integrity which serves as a negative feedback regulator of the transcriptional injury responses. *Smed-egr-5(RNAi)* (*early growth factor 5*), a crucial post-mitotic epidermal determinant, causes a loss of epithelial integrity and increased wound-induced gene expression (Tu et al., 2015). Similarly, *yki(RNAi)* animals had increased epidermal density concurrent with increased wound-induced gene expression (Figure 3A-D). In other systems, the roles for YAP in regulating cell density converge upon inputs such as cytoskeletal tension and mechanosensation (Dupont et al., 2011; Halder et al., 2012); yet, how tension may function in planarian biology, or in a regenerative context, remains unknown. In addition, other upstream cues may be other previously characterized pathways that have roles in regulating the planarian injury responses, including TGFβ signalling through the Activin-Follistatin axis, SMG-1 and mTORC, and JNK signalling (Almuedo-Castillo et al., 2014; Gaviño et al., 2013; González-Estévez et al., 2012). Yki/YAP also have described roles in interacting with each of these pathways, however, the interactions remain to be biochemically elucidated in planarians (Attisano and Wrana, 2013; Codelia et al., 2014; Csibi and Blenis, 2012; Sun and Irvine, 2013). Therefore, multiple wounding cues may converge on Yki to direct the injury responses. Taken together, we have shown that Yki is required to restrict the magnitude and duration of the injury responses, which coordinate proliferation to replace the missing tissue and patterning to integrate the newly differentiated cells; thus, Yki is essential in injury signalling that ultimately dictates scaling.

